# Unifying Gene Duplication, Loss, and Coalescence on Phylogenetic Networks

**DOI:** 10.1101/589655

**Authors:** Peng Du, Huw A. Ogilvie, Luay Nakhleh

## Abstract

Statistical methods were recently introduced for inferring phylogenetic networks under the multispecies network coalescent, thus accounting for both reticulation and incomplete lineage sorting. Two evolutionary processes that are ubiquitous across all three domains of life, but are not accounted for by those methods, are gene duplication and loss (GDL).

In this work, we devise a three-piece model—phylogenetic network, locus network, and gene tree—that unifies all the aforementioned processes into a single model of how genes evolve in the presence of ILS, GDL, and introgression within the branches of a phylogenetic network. To illustrate the power of this model, we develop an algorithm for estimating the parameters of a phylogenetic network topology under this unified model. The algorithm consists of a set of moves that allow for stochastic search through the parameter space. The challenges with developing such moves stem from the intricate dependencies among the three pieces of the model. We demonstrate the application of the model and the accuracy of the algorithm on simulated as well as biological data.

Our work adds to the biologist’s toolbox of methods for phylogenomic inference by accounting for more complex evolutionary processes.

## 1 Introduction

Independently evolving lineages of eukaryotic organisms are typically referred to as *species* (they may also be referred to as *populations* depending on the context and operational definition of those terms). Over evolutionary time scales, species lineages bifurcate to form two descendant species from a single ancestral species. This gives rise to a *species tree*, which is a phylogenetic tree describing the evolutionary history of a set of species. Each branch represents a species, and each bifurcation is represented by a *speciation* vertex. Rooted species trees are directed and acyclic.

A particularly powerful form of species tree in terms of its informativeness and utility for further analysis is a species *time tree*, which is a rooted species tree where each node has a height attribute which encodes the age of that node on a relative time scale (the expected number of mutations which have occurred per site in the genome) or an absolute time scale (years).

Estimating a species tree is challenging as gene trees are expected to be discordant with the species tree because of several well known processes. In describing those processes we will follow the convention of treating gene trees as evolving backwards in time (beginning in the present and ending in the past), and treating species trees (and later species networks, locus trees and locus networks) as evolving forward in time. The first process leading to discordance is incomplete lineage sorting (ILS), where multiple versions or *alleles* of a gene persist in a species up through to its ancestral species [9]. The second is horizontal gene transfer (HGT) through hybrid speciation [11], introgression [10], and speciation with gene flow [13]. This can lead to gene coalescent times which are younger than the earliest speciation event separating the corresponding species. The third is gene duplication and loss (GDL), where new copies of a gene are created at new loci in the genome, so that the relationship between sequences from different species at different loci (paralogs) reflects the duplication and loss process rather than the speciation process [4].

ILS has been addressed by years of research into the multispecies coalescent (MSC), a mathematical model which describes the evolution of gene trees within a species tree and naturally accommodates ILS [4]. In the MSC, the relationship between sequences from different species at orthologous (as opposed to paralogous) loci is represented by a gene tree, evolving within a species tree, and constrained so that its coalescent times must be older than the corresponding most recent common ancestors (MRCAs). A corresponding MRCA is the youngest speciation vertex whose leaf set contains all of the species which map to the leaf set of the gene subtree defined by a coalescent event.

More recently, HGT has been addressed by generalizing the MSC model to the multispecies network coalescent (MSNC) model, which represents the evolutionary history of species as a phylogenetic network [26]. Like their tree counterparts, rooted species networks are still directed and acyclic, but allow for two species to merge into a single lineage at a *reticulation* vertex. Each reticulation vertex has an inheritance probability *γ* ∈ [0, 1] which describes the proportion of genetic material of the child species that comes from one of the parent species. The proportion derived from the other parent species is 1 - γ. This flexible model of reticulate evolution can naturally accommodate hybrid speciation [30] and introgression [25]. Implementations of this method include `mcmc seq` in PhyloNet [24, 27] and SpeciesNetwork in BEAST [30].

GDL has been addressed by the development of models which add a third layer to the MSC between the species tree and the gene trees. This is known as the locus tree, and it contains vertices encoding duplication events, as well as vertices which directly correspond to the speciation vertices of the species tree [17]. The duplicate copy of a gene is assumed to reside in a new unlinked locus, so that there are multiple copies of a gene present in a single genome. The leaves of a single locus tree can therefore represent multiple loci, and the source of data in this model may be more appropriately termed “gene families” (cf. “genes”).

There is a one-to-many relationship between the species tree and locus tree speciation vertices, because original and duplicate subtrees will both be attached to a duplication vertex. Both original and duplicate subtrees contain all of the speciation vertices below the duplication (subject to losses), so there can be as many corresponding speciation vertices in the locus tree as there are duplication events above that speciation vertex.

DLCoal, the original implementation of the three-layer model [17], is relatively inflexible. It takes as input a gene tree topology, a species tree fixed in topology, branch lengths and effective population sizes, and rates of gene duplication and loss. From such input data it can estimate the locus tree, the mapping of gene tree coalescent vertices to locus tree branches, and the mapping of speciation vertices in the locus tree to the species tree. DLCoal also relies on the accuracy of the supplied gene tree topology, which may contain errors due to the gene tree inference method or insufficient information in the original multiple sequence alignment (MSA). A later method, DLC-Coestimation [29], avoids that potential issue by jointly estimating the gene tree along with the locus tree and reconciliations and mapping directly from a gene family MSA.

The most recent implementation of the three-layer model jointly estimates the species, locus and gene tree topology and times, as well as general parameters including duplication and loss rates from the MSAs of multiple gene families [6]. In a simulation study, this method was able to successfully infer the species tree topology, and outperformed using the MSC model alone (without accounting for GDL) when estimating species divergence times [6].

While the above methods either account for both ILS and HGT, or for both ILS and GDL, no model has been designed or implemented that account for all three processes which generate gene tree discordance; ILS, HGT and GDL. Here we present a new model which extends the MSNC to a three-layer model by adding a locus network between the species network and gene trees. This new model accounts for HGT at the species network level, GDL at the locus network level, and ILS at the gene tree level. We have implemented a maximum *a posteriori* (MAP) search for this model which jointly estimates the speciation times, inheritance probabilities and duplication and loss rates. Using simulation experiments, we show that it can accurately infer the aforementioned parameters.

We also used simulated data and an empirical data set of six yeast species to study the difference in accuracy between our new method and an MSNC method which does not account for GDL. Results from those experiments showed that accounting for GDL in addition to ILS and HGT is particularly important when estimating reticulation times.

## 2 Methods

Similar to the three-layer model of [17], we develop a three-layer model that uses a locus network (different from the locus tree of [17]) as an intermediate layer between the species network and gene tree. This structure allows for unified modeling of coalescence and GDL, where all coalescence events are captured by the relationship between the gene tree and locus network, and all GDL events are captured by the relationship between the locus network and phylogenetic network. The reticulation events (e.g., introgression) are captured by the fact that the species and locus structures are both networks, rather than trees.

### 2.1 The Three-Layer Model

A species network 𝕊 = (*V* (*S*), *E*(*S*), *τ*^*S*^) is a directed acyclic graph depicting the reticulate evolutionary histories of a set of species where *V* (*S*) is the set of vertices in the network, *E*(*S*) is the set of edges and *τ*^*S*^ contains the set of branch lengths of the edges. We use *S* to denote *{V* (*S*), *E*(*S*)}. Further, *V* = *r* ∪ *V*_*L*_ ∪ *V*_*T*_ ∪ *V*_*N*_ where *r* is the root of the network, *V*_*L*_ is the set of leaf vertices, *V*_*T*_ denotes the set of tree vertices with two children and one parent and *V*_*N*_ represents the set of reticulation vertices with one child and two parents. The set of all internal vertices is *IV* (*S*) = *r* ∪ *V*_*T*_ ∪ *V*_*N*_. If *u* has only one parent, we call this parent *pa*(*u*). The set of children of *u* is denoted as *c*(*u*). For each reticulation vertex *u* with two parents *v* and *w*, there is an inheritance probability *γ ∈* [0, 1] such that the probability of locus *u* inheriting from *v* is *γ* and inheriting from *w* is 1 − *γ*. *Γ* is a vector of all inheritance probabilities for all vertices in *V*_*N*_, *Γ* ((*v, u*)) = *γ* and *Γ* ((*w, v*)) = 1 − *γ*. The population sizes are denoted as *N*^*S*^ and the population size on branch *e*(*u, v*) is *N*^*S*^((*u, v*)).

#### Locus Networks and Locus-network-to-species-network Reconciliation

A locus network 𝕃 = (*V* (*L*), *E*(*L*), *τ* ^*L*^) is generated by applying duplication and loss events onto the species network with a top-down birth-death process [2, 1, 21]. Birth events create new loci by duplicating an existing locus, and death (loss) events eliminate loci so that it will have no sampled descendants. Fully describing the result of this process requires a reconciliation *R*^*L*^ from the locus network to the species network, where the vertices on the locus network can be mapped to either the vertices or the branches of the species network. If *u ∈ V* (*L*) is mapped to a species network vertex, then we call it a speciation vertex; the set of speciation vertices is denoted as *V*_*S*_ (*L*). If it is mapped to a species network branch; we call it a duplication vertex and the set of duplication vertices is denoted as *V*_*D*_(*L*). Branches with no existing leaf vertices are pruned out (Figure 1). For a duplication, a new locus is generated, so a mapping *δ*(*u, v*) = 1 or *δ*(*u, v*) = 0 is used to indicate whether (*u, v*) leads to the new (daughter) locus or if (*u, v*) is the mother branch where *u* is the duplication vertex. The population size of branch *e* = (*u, v*) in the locus network is the population size of the branch *e*′ = (*w, x*) on the species network where *R*^*L*^(*u*) = (*w, x*) or *R*^*L*^(*v*) = (*w, x*) or *R*^*L*^(*u*) = *w, R*^*L*^(*v*) = *x*. Similarly, *Γ* ((*u, v*)) = *Γ* ((*w, x*)) where (*w, x*) *∈ E*(*S*) if *R*^*L*^(*u*) = (*w, x*) or *R*^*L*^(*v*) = (*w, x*) or *R*^*L*^(*u*) = *w, R*^*L*^(*v*) = *x*. It is important to note that reticulation edges present in the species network may be deleted from the locus network (as in the (*F, X*) branch leading to the B2 locus in Figure 1), so the locus network can be a tree or more tree-like with fewer reticulation vertices than the species network.

**Fig. 1.**
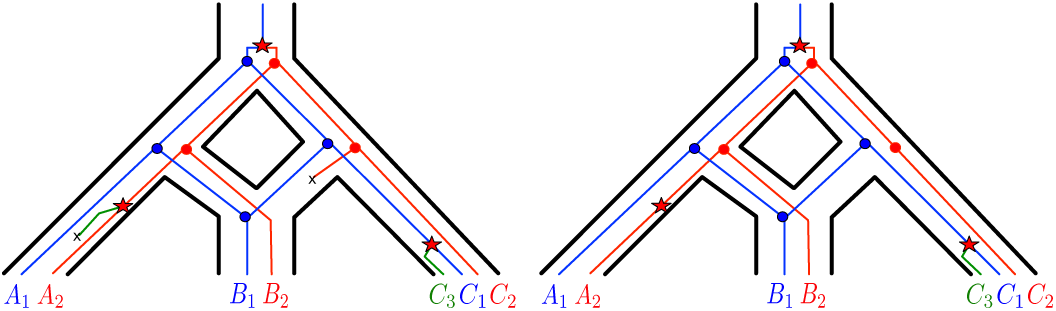
A gene duplication and loss scenario inside of a species network on three species *A, B*, and *C*. (Left) The complete locus network embedded in the species network, produced by a birth-death process, and containing all duplication and loss events. (Right) Lineages in the locus network with no sampled loci due to loss events are pruned from the locus network, resulting in the observed locus network. Extinct lineages are deleted. Duplication, loss, and speciation/hybridization events are represented by ⋆, ×, and *•*, respectively. New lineages arising from duplication are colored red and green.

#### Gene Trees and Gene-tree-to-locus-network Reconciliation

A gene tree 𝔾 = (*V* (*G*), *E*(*G*), *τ* ^*G*^) describes the evolution of lineages and the definitions of vertices are similar with those in the species network and locus network. The reconciliation from the gene tree to the locus network is denoted by *R*^*G*^. The two reconciliations *R*^*L*^ and *R*^*G*^ are collectively denoted by *R*. For each locus network branch *e* = (*u, w*) with *δ*((*u, w*)) = 1, the coalescent time of every gene vertex mapped to the leaf vertices under *w* must be more recent than *u*. Also, we define *M* as the mapping from the gene tree leaf vertex-set to the locus network leaf vertex-set. *M* indicates what gene is from what locus in the locus network. Figure 2 shows the reconciliations from the gene tree to the locus network (*R*^*G*^) and from the locus network to the species network (*R*^*L*^).

**Fig. 2.**
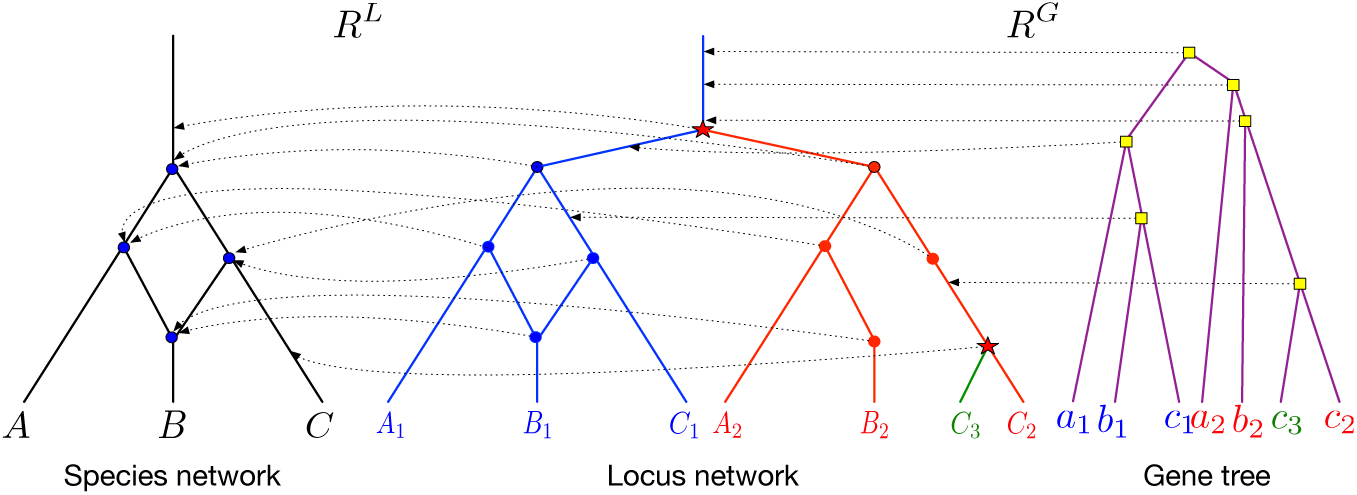
Gene duplication/loss events are obtained by mapping the nodes of the locus network onto the branches of the species network, via reconciliation *R*^*L*^ (the dotted arrows from the locus network to the species network). Coalescence events are obtained by mapping the gene tree nodes onto the branches of the locus network, via reconciliation *R*^*G*^ (the dotted arrows from the gene tree to the locus network). Duplication, loss, speciation/hybridization, and coalescence events are represented by ⋆, ×, ●, and ■, respectively.

### 2.2 Model Assumptions

In this model, we need to make some assumptions as made in [17, 29, 6].

1. After the duplication, the daughter locus becomes totally unlinked and any further evolution of the mother and daughter loci, as well as the coalescent histories of the mother and daughter genes, are independent conditional on the species network topology, times and population sizes. Thus we can calculate the coalescent probabilities separately for each locus, and use the product as the gene family coalescent probability.
2. At a locus level, hemiplasy [3] is assumed to be non existent in this model. In other words, for each duplication and loss event, the resulting addition or deletion of locus will be transmitted universally to all descendent species. This allows us to explain all unobserved loci by means of gene loss.
3. In our present implementation, one individual per species is sampled for each locus. For pure multi-species coalescent models, e.g. [22, 14, 20, 5, 28], it is often the practice to allow for multiple individuals for each species. However in the context of modeling gene duplication / loss and coalescence, allowing multiple individuals might cause problems, please refer to [19] for a study to investigate the feasibility of reconciliation with multiple individuals under duplication / loss and coalescence model.

### 2.3 Probability Distribution

For a species network 𝕊 and a set of gene families 𝔾𝔽 with each member 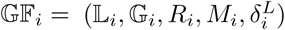, and parameters *θ*, the posterior *p*(𝕊, 𝔾𝔽, *θ*| *D*) given observed DNA sequences *D* is

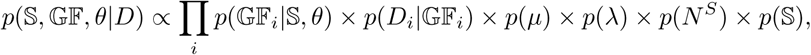

where *D*_*i*_ is the DNA sequences for 𝔾𝔽 _*i*_ and *θ* = {*µ, λ, ψ, Γ, N*^*S*^} which are the duplication rate, loss rate, substitution rate, inheritance probabilities and population size respectively. The term *p*(𝔾𝔽_*i*_|𝕊, *θ*) can be decomposed into (we drop the subscript *i* for readability) the product *p*(*G, τ* ^*G*^, *R*^*G*^ |*L, τ* ^*L*^, *δ*^*L*^, *M, Γ, N*^*S*^) *× p*(*M*|*L, τ* ^*L*^, *R*^*L*^, *δ*^*L*^) *×p*(*L, τ* ^*L*^, *R*^*L*^, *δ*^*L*^ |*S, τ* ^*S*^, *µ, λ*), and we have *p*(*D*|𝔾𝔽_*i*_) = *p*(*D* |*G, τ* ^*G*^). The term *p*(*G, τ* ^*G*^, *R*^*G*^ |*L, τ* ^*L*^, *δ*^*L*^, *M, Γ, N*^*S*^) is the probability of the gene tree coalescing in the locus network under a bounded coalescence model where gene lineages originated from gene duplication events must coalesce earlier than the duplication event. The bounded coalescence model is extended from [17] and gains the capacity to handle hybridization events. The details are in Supplementary Materials.

The term *p*(*M* |*L, τ* ^*L*^, *R*^*L*^, *δ*^*L*^) is the probability of the map of gene tree leaves to locus network leaves. Since we assume no prior knowledge of locus information of each sampled gene copy from a certain species, the mapping has a uniform distribution based on the number of possible permutations:

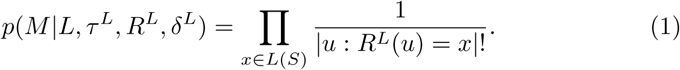

The number of permutations is constant for a given data set *D*, so for identification of the MAP topology or algorithms like MCMC which use unnormalized posterior probabilities not scaled by 1*/P* (*D*), the calculation of this prior is unnecessary. The term *p*(*L, τ* ^*L*^, *R*^*L*^, *δ*^*L*^ |*S, τ* ^*S*^, *µ, λ*) is the probability of the locus network generated inside of the species network with duplication rate *µ* and loss rate *λ* and is also derived in Supplementary Materials. The term *p*(𝕊) is the prior of the species network which is a compound prior with uniform prior on the topology and exponential prior on divergence times as in [7, 23].

### 2.4 MAP Inference of the Parameters of a Fixed Network Topology

Our goal is to find the maximum *a posteriori* (MAP) estimate of the parameters; that is,

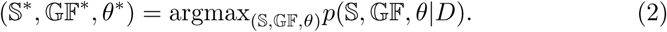

In this present work, we will focus on inferring the species network parameters— times, population sizes and inheritance probabilities—with the topology being fixed, as well as locus networks, gene trees and reconciliations between them. General parameters such as duplication and loss rates are also inferred. Because of the hierarchical nature of the generative model, changes on higher level components will influence lower level components as well. For example, changing the heights of the species network vertices will also change the heights of corresponding locus network vertices. We developed four groups of operators, each working on different levels of the model, which we describe in detail in Supplementary Materials due to space limitation. The first group makes changes to the species network and can also alter the locus networks and gene trees. The second group changes the locus network and can also alter the gene trees. The third group makes changes to the gene trees alone, while the fourth applies to the macroevolutionary rates, which in our implementation is limited to the duplication and loss rates.

### 2.5 Results and Discussion

### 2.6 Performance on Simulated Data

#### Simulation Setup

We simulated DNA sequence data for multiple gene families with our gene tree simulator and Seq-Gen [15]. Our gene tree simulator employs the hybridization-duplication-loss-coalescence model and operates in two phases. First, it generates the locus network within a predefined species network by simulating duplications and losses. Then the gene tree is simulated under a coalescence model along the locus network. If the gene lineages could not coalesce before the duplication event backward in time, it will be rejected and retried until it can or 10^8^ trials are done where the locus network will be rejected and regenerated. Locus network with less than 3 species left will be rejected. Once the gene trees were generated, the program Seq-Gen [15] was used to simulate the evolution of DNA sequences down the gene trees under a specified model of evolution. In all simulations reported here, we used the Jukes-Cantor model of evolution [8] to generate 1000 bp long DNA sequences. For Experiments 1 to 3, we used the network of Figure 3 as the model species network. Population sizes are given as the number of diploid individuals, and specified duplication/loss rates and population sizes were set to be the same across all branches of the model networks. A mutation rate of 10^−9^ was used for all simulation experiments.

**Fig. 3.**
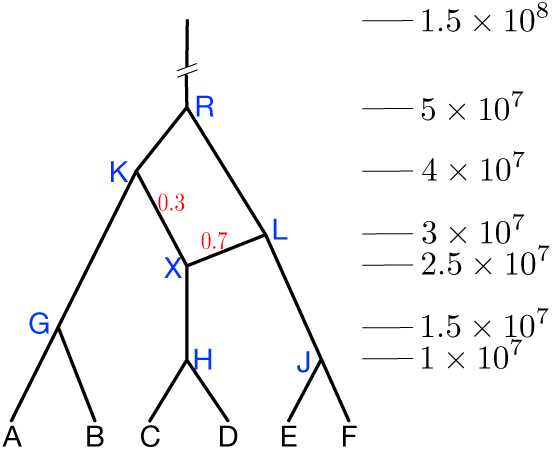
The model network used to simulate data for Experiments 1 to 3. The values on the right correspond to divergence times of the nodes in number of generations. The inheritance probability values are shown on the reticulation edges.

#### Experiment 1: Testing the Effects of GDL Rate and Population Size

In this experiment, different settings of duplication/loss rates and population sizes were used to test how these parameters would affect the accuracy of inferences. The duplication and loss rates (both were equal) used were 5 × 10^−10^, 10^−9^ and 2.5 × 10^−9^ and the population sizes were 10^6^, 4 × 10^6^, and 8 × 10^6^. For each of the 9 different settings of duplication/loss rates and population size, we generated 10 replica each with 50 gene families and ran 15 million iterations for each data set. First, we calculated the average difference of the estimated divergence times and the true values in unit of population mutation rates for the 9 settings based on the 10 replica for each setting. Most estimates of the divergence times are accurate across different settings, with the exception of the reticulation time “X” which appears to be less identifiable at smaller population sizes (Figure 4). We calculated the difference between the estimated inheritance probabilities and the true value (0.3) on (*K, X*) (Figure 5). No consistent trend was observed in the accuracy of inheritance probability estimates over the ranges of population size and duplication and loss rates studied. To assess the accuracy of locus networks and gene trees, we calculated the topological error in metrics developed by [12] between the estimated locus networks and RF distance between gene trees [18] and the true networks and trees. Overall our method shows very good accuracy (indicated by topological distances close to 0). The average distance for the locus networks increases as the duplication and loss rate increases, but it appears invariant to different population size. This makes sense because the locus networks are determined by the duplication and loss events not the ILS events. If ILS, GDL and HGT are absent all gene tree topologies will identical to the species tree topology, and be perfectly accurate when the species tree topology is fixed at the truth. However in our model gene trees can vary because of all three processes. The prevalence of ILS is partly dependent on population sizes, and therefore it is unsurprising that we show gene tree topological error consistently increasing as population sizes get larger (Figure 6). Finally, we assessed the method’s performance in terms of estimating the duplication and loss rates. As the results show, the method performs well at estimating both rates under the range of population sizes and duplication and loss rates studied (Figure 7).

**Fig. 4.**
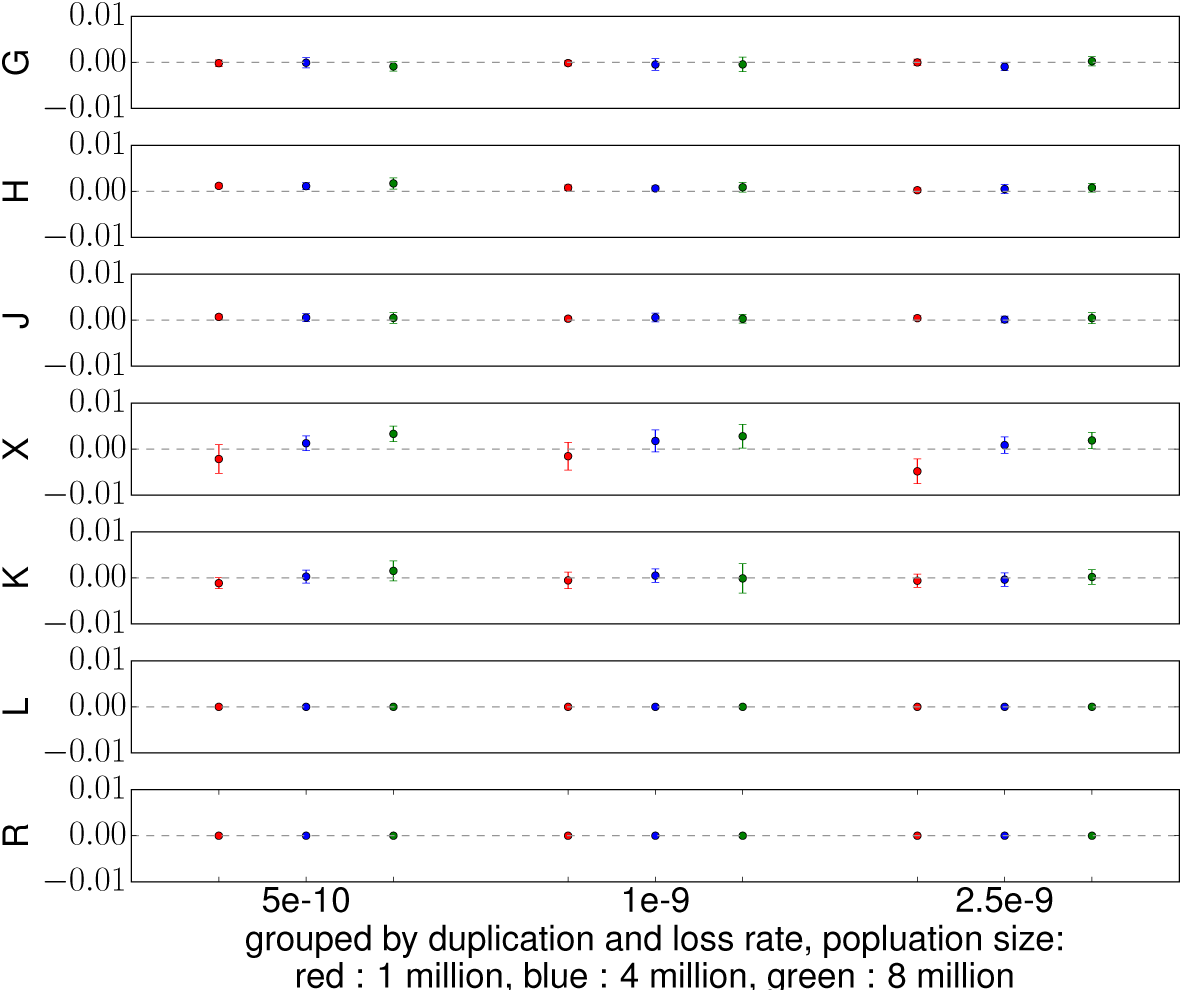
The average difference of the estimated divergence times and the true values. Divergence times for each panel from top to bottom are for vertices: *G, H, J X, K, L* and *R* respectively. In each panel, estimations are grouped by duplication and loss rates and the the red, blue and green color correspond to different population size. Standard deviation is represented as vertical bar.

**Fig. 5.**
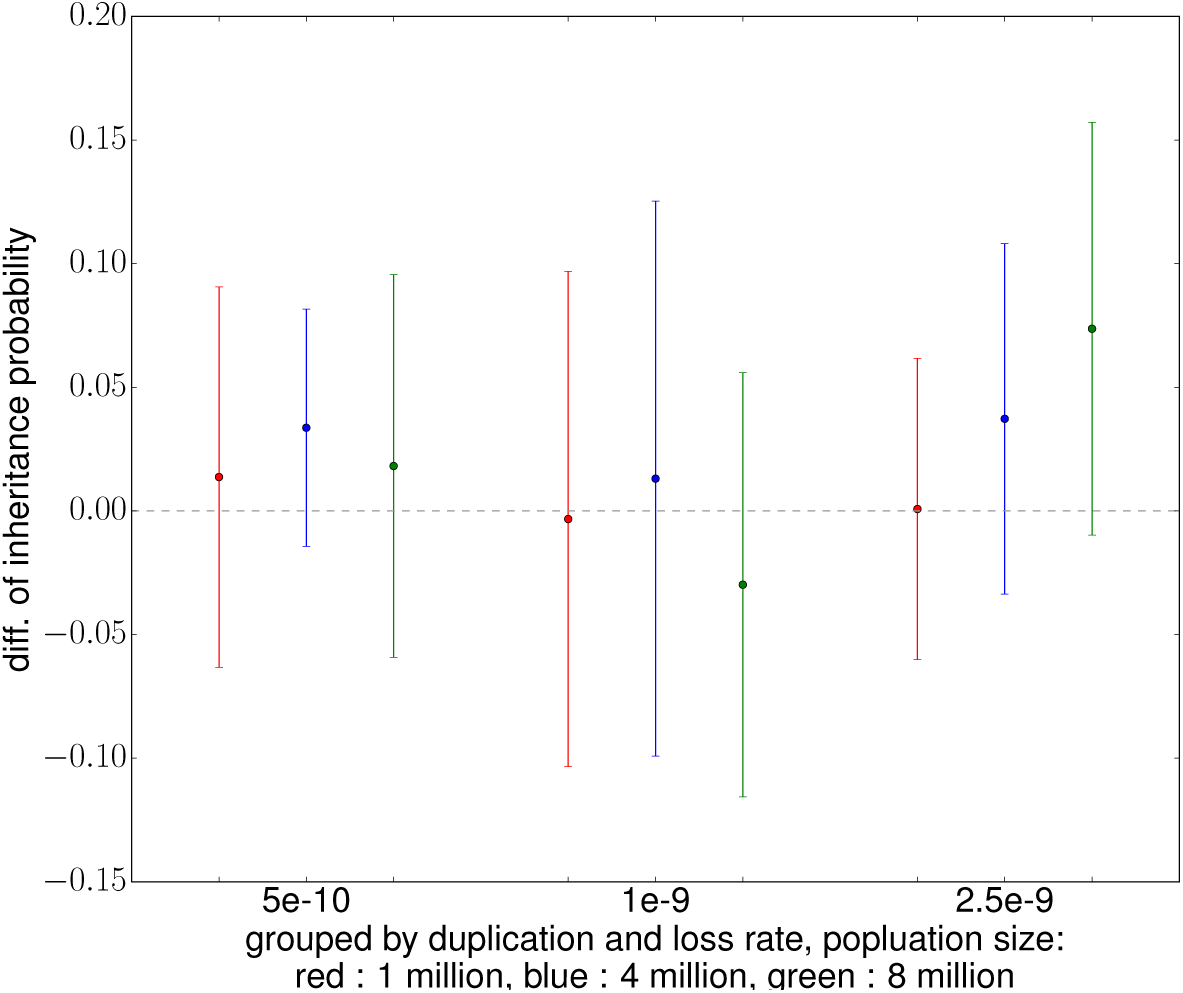
The average difference between the estimated inheritance probability on branch (*K, X*). Estimation are grouped by duplication and loss rate used and the the red, blue and green color correspond to different population size. Standard deviation is represented as vertical bar.

**Fig. 6.**
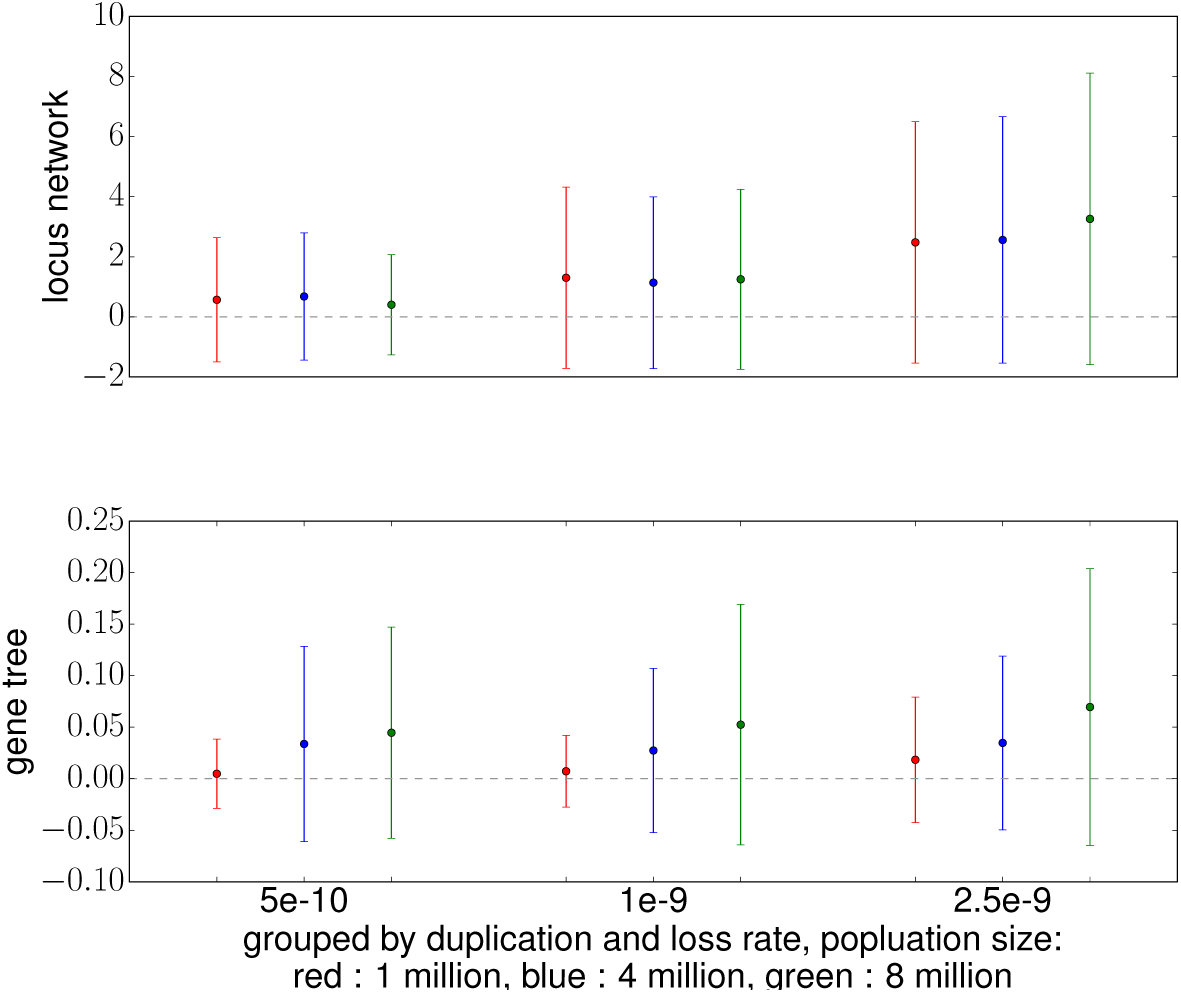
The average topological distances between the estimated networks / trees and its corresponding true networks /trees. Values for the locus networks and gene trees are shown in the top and bottom panel respectively. In each panel, estimations are grouped by duplication and loss rates and the the red, blue and green color correspond to different population size. Standard deviation is represented as vertical bar.

**Fig. 7.**
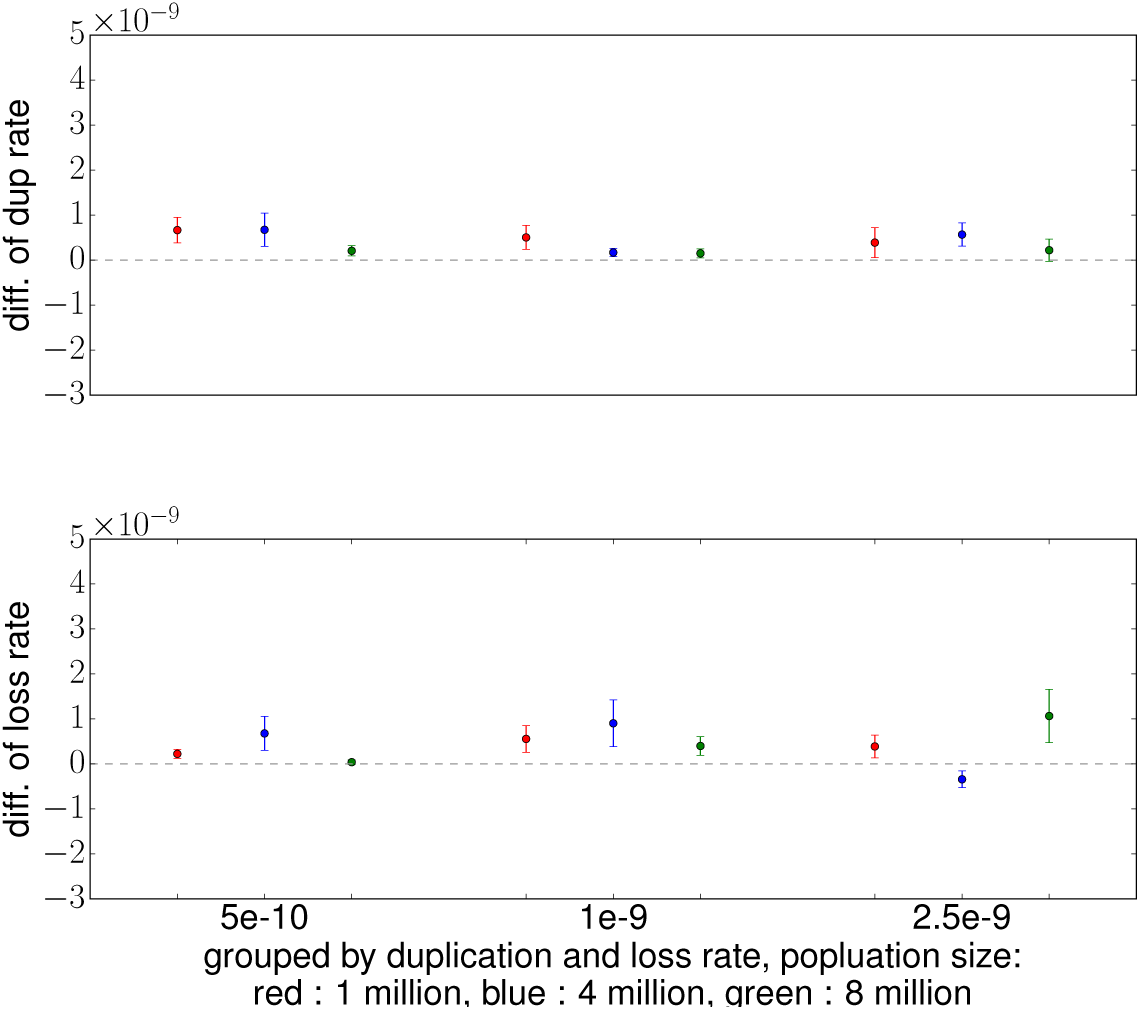
The average difference of estimated duplication and loss rates from true ones for each setting. Estimations for the duplication and loss rates are shown in the top and bottom panel respectively. In each panel, estimations are grouped by duplication and loss rates and the the red, blue and green color correspond to different population size. Standard deviation is represented as vertical bar.

#### Experiment 2: Testing the Effect of the Number of Gene Families

In order to determine how our method performs given larger or smaller data sets, we varied the number of gene families (5, 10, 25, and 50) under one setting of duplication/loss rate (2.5 × 10^−9^) and population size (4 × 10^6^). 10 replica for each number of gene families were simulated and 15 million iterations were run for each data set. Results (Figure 8) show that even for 5 gene families, a relatively small number, the estimated divergence times are generally accurate especially for *H* and *R*. The accuracy and precision improve as more gene families are used for example for node *G, J* and *L*.

**Fig. 8.**
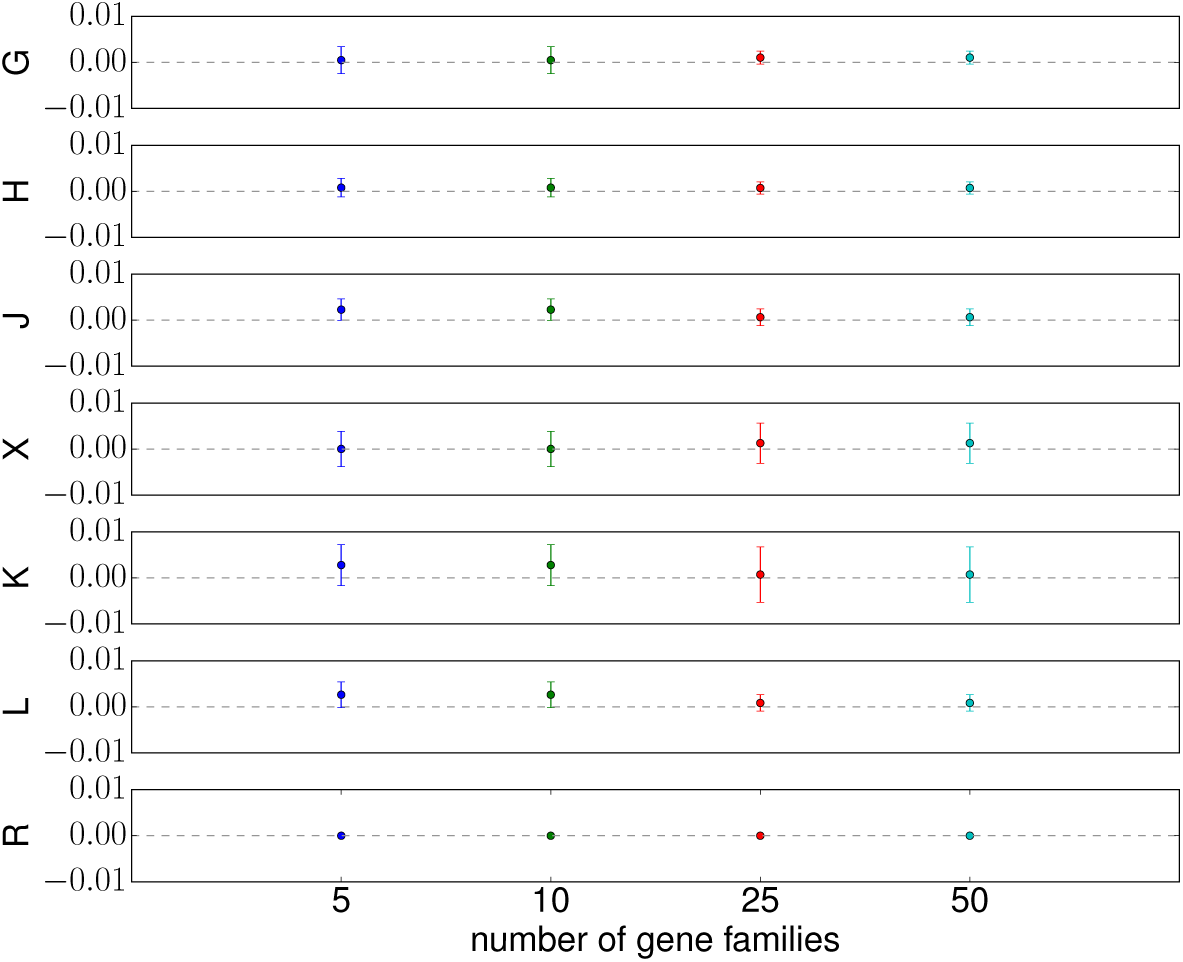
The difference between the estimated divergence times and the true values for 4 different settings of number of gene families (5, 10, 25, 50) for internal vertices. Divergence times for each panel from top to bottom are for vertices: *G, H, J X, K, L* and *R* respectively. Standard deviation is represented as vertical bar.

We tested the inference of other parameters. Figure 9(a) shows that both the accuracy and precision of the inheritance probability improved for larger numbers of gene families. The accuracy of the duplication rate both appear to improve slightly with more data. The loss rate, while accurately estimated, did not show any consistent trends. As the results show, the accuracy of the locus networks and of the gene trees seems to be stable across different settings in terms of both mean and standard deviation. As gene tree topologies, while independent, are conditioned on the species network topology, when the species network topology is fixed even without any data there will already be a lot of information in the model on the gene tree topologies. Also, the locus networks are independent for each gene tree conditioned on the species network topology. So increasing the number of gene trees will not improve the overall accuracy of gene tree estimates to the same extant as when jointly estimated with the species tree or network topology (Figure 9(b)).

**Fig. 9.**
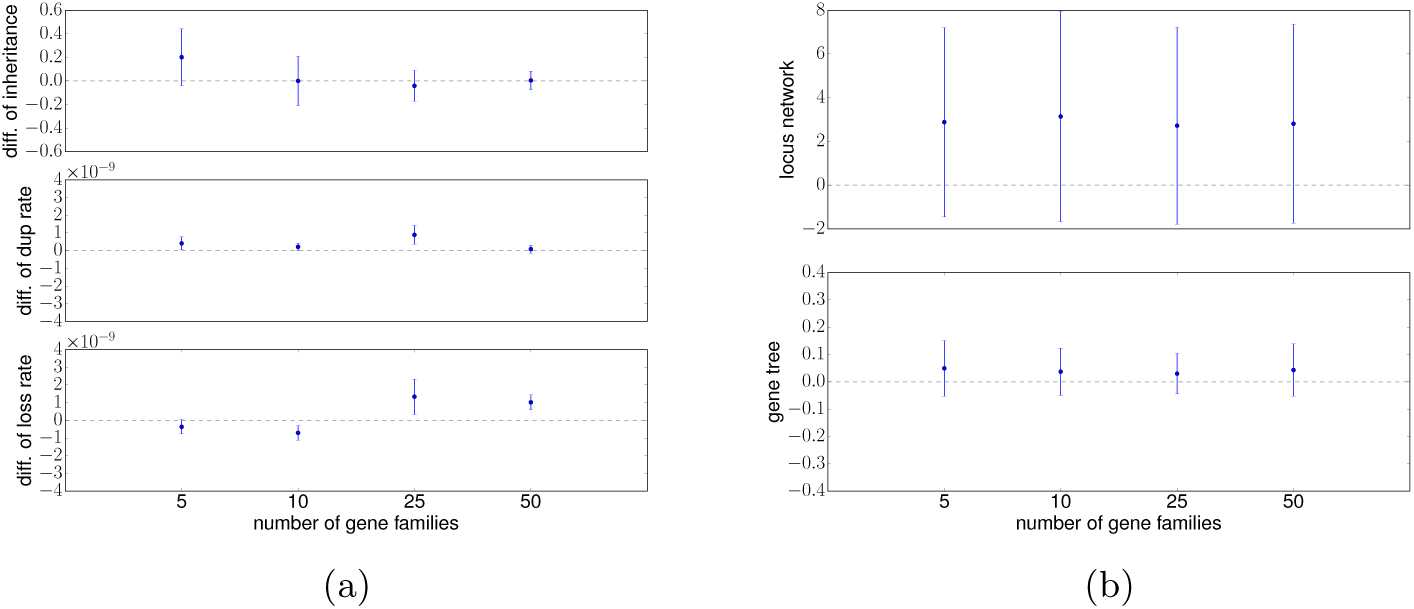
(a) The difference of estimated parameters from the true values. Top: difference between estimated inheritance probability and the true value (0.3) on (*H, X*). Middle: difference of estimated duplication rate and true value. Bottom: difference of estimated loss rate and true value. The number of gene families used as input to the inference method is shown on the x-axis. Standard deviation is represented as vertical bar. (b) The average topological distances between the inferred and true networks or trees. Top: Locus network difference. Bottom: Gene tree difference. The number of gene families used as input to the inference method is shown on the x-axis. Standard deviation is represented as vertical bar.

#### Experiment 3: Comparing Inference With and Without GDL

In this experiment we set out to test how a method that accounts only for incomplete lineage sorting but ignores duplication and loss would perform as compared to our model here. To achieve this, we ran our method and a Bayesian MCMC species network inference method (the `mcmc seq` command, with the species network fixed, and using the MAP estimation) in PhyloNet [27] which implements the method of [24]. We simulated 10 replica under duplication and loss rates of 2.5 × 10^−9^ and population size 10^7^ and 50 gene families for each data set. For each gene family we randomly selected one gene copy for each species if there was at least one. As a result, around half of the sequences in the gene families were kept after this pruning of the data sets. We fed the sequences to both methods and ran them both for 15 million iterations. Figure 10 shows the divergence time estimates obtained by the two methods for the six internal vertices of the model species network. Our results show that our method, which accounts for gene duplication and loss even with a single sampled locus per species, more accurately estimated speciation and reticulation times. This was particularly true of the reticulation vertex, where `mcmc seq` dramatically underestimated the reticulation time (Figure 10). Also, we have a better estimation of the inheritance probabilities than `mcmc seq`. Our estimation is 0.268 and `mcmc seq` had estimation of 0.464 where the true value is 0.3.

**Fig. 10.**
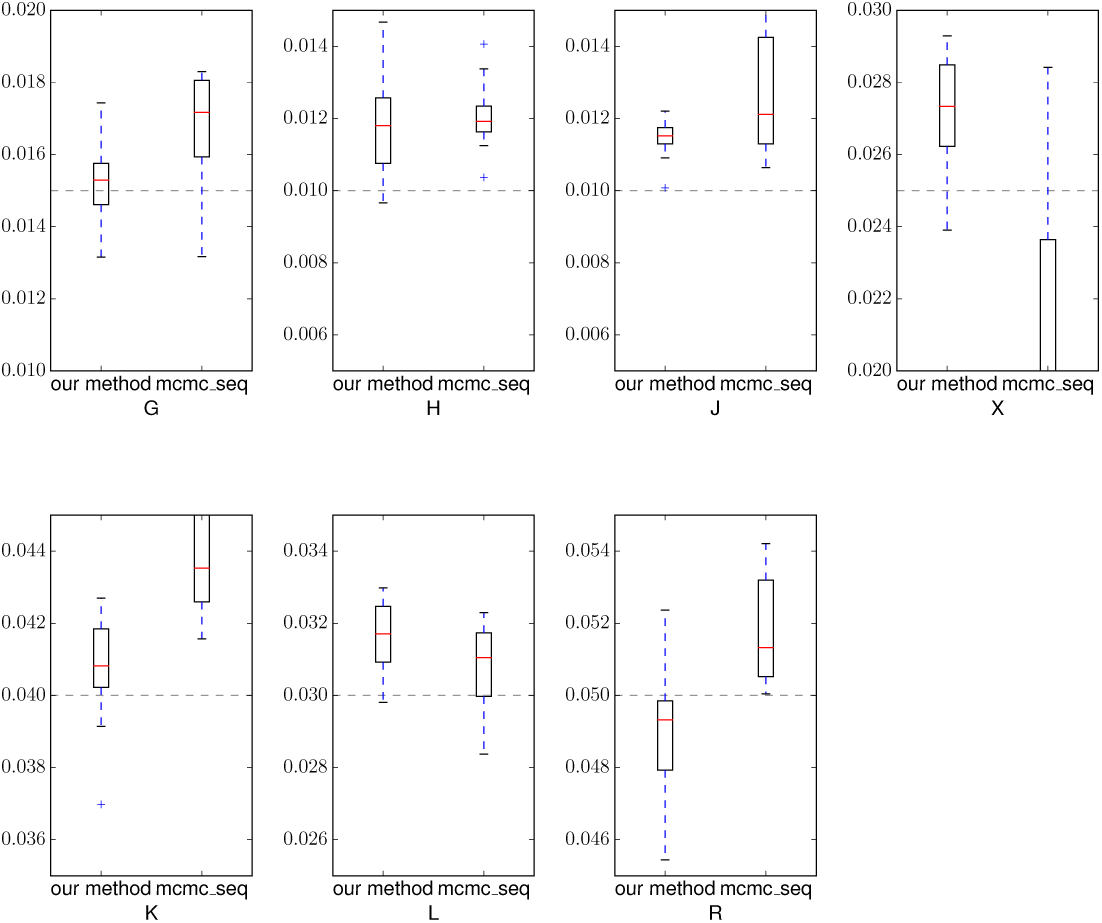
The divergence times in population mutation rates for the ancestors of *G, H, J X, K, L* and *R* respectively estimated by our method (red) and the species network inference method in `mcmc seq` (orange). Vertical bars are the true divergence times.

### 2.7 Biological Data

We used the yeast genome data set with duplications reported on http://compbio.mit.edu/dlcoal/ and randomly selected two data sets restricted to six genomes. One consists of 100 gene families each with exactly one copy for each species with alignments 1000nt–2000nt in length; the other consists of 100 gene families with possibly multiple or 0 copies for each species with alignments 1000nt–2000nt in length. We used 10^−10^ as duplication and loss rate and 4 × 10^−10^ as mutation rate and 10^7^ as population size which are comparable with the settings used in [16]. Then we fed `mcmc seq` with the first data set and ran the command 10 times for 15 million iterations each with the maximum number of reticulation vertex set to be one. The most prevalent topology is shown in Figure 11(a) and appeared in 7 of the 10 runs. We then fed our method with the second data set and run 7 times each with 15 million iterations. A table of the average estimated divergence times is given in Figures 11(b). We can see that most of the branch lengths are similar and the only significant differences are at the branch lengths of vertices J, X and L. Given our method is better at estimating divergence times given results from Experiment 3, it seems that the ones obtained by our method here are probably more accurate estimations.

**Fig. 11.**
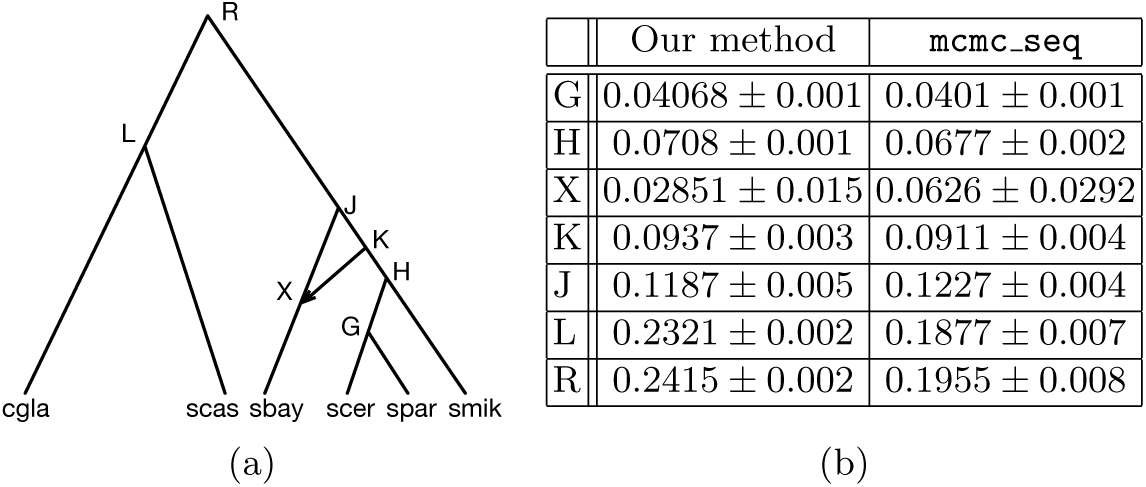
(a) The yeast species network topology inferred by `mcmc seq` on the 100 gene families. (b) Mean and standard deviation of the estimated divergence times of `mcmc seq` and our method (std’s smaller than 0.001 are rounded to 0.001).

Under the multispecies (network) coalescent model without duplication and loss, the most probable explanation for a lack of coalescence between contem-poraneous gene lineages is if they are embedded in separate species branches. For example, they may be embedded in different parent branches of a reticulation vertex. This aids in estimating the timing of reticulation vertices by setting their height to around where the coalescent rate of embedded lineages traversing that reticulation cease declines. However when duplication and loss exists, the gene lineages may be embedded in the same species network branch but different locus network branches due to a duplication event. Therefore reticulation times may be underestimated when not accounting for duplication and loss, as a lack of coalescence may be attributed to underestimated reticulation vertex heights. This phenomenon can explain the more recent reticulation times estimated by `mcmc seq`, known with certainty to be underestimated in the case of the simulated data.

The inheritance probability on branch (*R, X*) estimated by `mcmc seq` was 0.503 ± 0.147 while the value estimated by our method was 0.461 ± 0.091. The error is the standard deviation among runs, and shows that the estimated inheritance probabilities of the two methods are very close.

## 3 Conclusions

The availability of genome-wide data from many species and, in many cases, from several individuals within species, promises to significantly improve the accuracy of inferred evolutionary histories and estimated evolutionary parameters. However, along with this promise comes the major challenge of accounting for the various evolutionary processes that act simultaneously within and across species boundaries to create genomic mosaics of different evolutionary pathways. These processes include incomplete lineage sorting, hybridization and introgression, gene duplication and loss, and recombination.

The mathematical and computational communities are hard at work refining existing models and developing new ones to increasingly account for the complexities of evolution. Most relevant to our work here is that recent development of statistical methods for phylogenetic network inference. Those methods employ the multispecies network coalescent to account simultaneously for incomplete lineage sorting and introgression as the sources of heterogeneity across genomic loci. However, these methods do not account for even more ubiquitous processes that have been documented across all branches of the Tree of Life, namely gene duplication and loss.

In this work, we developed a probabilistic model that simultaneously accounts for hybridization, gene duplication, loss and ILS. We also devised a stochastic search algorithm for parameterizing phylogenetic networks based on this model. This algorithm provides estimates of evolutionary parameters, as well as gene histories and their reconciliations. Results based on simulation studies show good performance of the algorithm as well as insights obtained by employing the new model as compared with existing models that exclude gene duplication and loss.

We identify three natural directions for future research. First, while in this work we assumed a fixed phylogenetic network topology, in most empirical studies such a topology is not given or known. Developing a method that infers the phylogenetic network, along with all the parameters that the current method estimates, is essential for proper application of the model. Second, while this work focused on obtaining point estimates of the phylogenetic network’s parameters, developing a method that estimates a posterior distribution on the space of phylogenetic networks and their parameters would provide additional information, including assessment of statistical significance and the uniqueness and distinguishability of optimal solutions. Third, the computational bottleneck in this domain stems from the time it takes to compute the likelihood of a given point in the parameter space as well as from the need to walk an enormous and complex space of such parameters. For example, it took between 15 and 20 hours for a single run of 15 million iterations on a data set with four or five species and 50 gene families. Developing algorithmic techniques and potentially alternative likelihood functions to speed up these calculations is imperative for this work to be applicable to data sets of the scale that biologists can now generate using the latest sequencing technologies.

