## Supplementary Materials for "Unifying Gene Duplication, Loss, and Coalescence on Phylogenetic Networks"

### 1 Probability Distribution

The term $p(G, \tau^G, R^G|L, \tau^L, \delta^L, M, N^S)$ is the probability of the gene tree coalescing in the locus network under a bounded coalescence model where gene lineages originated from gene duplication events must coalesce earlier than the duplication event backward in time. The bounded coalescence model is extended from [5] and gain the capacity to handle hybridization events.

$$p(G, \tau^G, R^G|L, \tau^L, \delta^L, M, \Gamma, N^S) \tag{1}$$

$$= \frac{p_{CL}(G, \tau^G, R^G|L, \tau^L, \delta^L, M, \Gamma, N^S)}{\prod_{u \in V_D(L)} p_{BC}(\tau(G_u) < \tau(u)|L, \tau^L, M, \Gamma, N^S)} \tag{2}$$

where $G(u)$ is the root of sub gene tree reconciled under $u$ and the numerator of Equation 2 can be further decomposed as:

$$p_{CL}(G,\tau^G, R^G|L, \tau^L, \delta^L, M, \Gamma, N^S) = \tag{3}$$

$$p(G, R^G|L, \tau^L, \delta^L, M, \Gamma, N^S)\times \tag{4}$$

$$p(\tau^G|G, R^G, L, \tau^L, \delta^L, M, \Gamma, N^S) \tag{5}$$

Because in the model, the reconciliation only consider which gene tree vertex is mapped to which locus network branch, the embeddings of the exact path of the gene lineages are transparent. So, we borrowed the idea of Ancestral Configuration from [6] and developed algorithm to calculate the probability under the duplication, loss and ILS setting. as is defined in [3].
$p(L, \tau^L, \delta^L|S, \tau^S, \mu, \lambda)$ is evaluated as evolving gene duplication and loss events using a Birth-Death process inside of a species network. The probability of an event corresponding to a vertex on the locus network $u$ is denoted as $p_v(u)$. For $u \in V^L$ with only one child, the probability of losing the locus from $u$ downward to corresponding species subnetwork is denoted as $e(u)$. In this calculation, we made simplifying assumption that the species subnetwork is treated as a MUL

* This work was supported in part by NSF grants DBI-1355998, CCF-1302179, CCF-1514177, CCF-1800723, and DMS-1547433.

tree [7] presentation and duplication and loss on branches under hybridization are independent. The probability of no duplications or loss happens on a branch $(u, v)$ is denoted as $p_{nb}((u, v))$. We denote the probability of the locus network rooting at $r$ with single child $u$ as $p_{BD}(r, u, |S, t^S, \mu, \lambda)$

$$
\begin{aligned}
p(L, &\tau^L, R^L, \delta^L | S, \tau^S, \mu, \lambda) && (6)\\
&= p_{nb}((r, u)) \times p_v(u) \times && (7)\\
&\prod_{u}^{V-\{r\}} 1 \times \begin{cases} p_{nb}((u, v)) \times p_{nb}((u, w)) & \text{if } c(u) = \{v, w\} \\ p_{nb}((u, v)) \times e(u) & \text{if } c(u) = \{v\} \& R^L(u) \notin V_N(S) \\ p_{nb}((u, v)) & \text{if } c(u) = \{v\} \& R^L(u) \in V_N(S) \\ 1 & \text{if } c(u) = \{\} \end{cases} && (8)
\end{aligned}
$$

where

$$p_v(u) = \begin{cases} \mu \text{ if } u \in V_D(L) \\ 1 \text{ if } u \notin V_D(L) \end{cases} \tag{9}$$

and

$$p_{nb}((u, v)) = (1 - \mu - \lambda)^{\tau(u) - \tau(v)} \tag{10}$$

We define :

$$f_\tau = \frac{\mu(1 - e^{-(\mu-\lambda)^\tau})}{\mu - \lambda e^{-(\mu-\lambda)^\tau}} \tag{11}$$

$$g_\tau = \frac{\mu - \lambda}{\mu - \lambda e^{-(\mu-\lambda)^\tau}} \tag{12}$$

and

$$\lim_{\mu \to \lambda} f_\tau = \frac{\lambda \tau}{1 + \lambda \tau} \tag{13}$$

$$\lim_{\mu \to \lambda} g_\tau = \frac{1}{1 + \lambda \tau} \tag{14}$$

Further, we define $p_d(\tau)$ as

$$p_d(\tau) = \begin{cases} g_\tau (1 - f_\tau) f_\tau^{d-1}, & d > 0 \\ 1 - g_\tau, & d = 0 \end{cases} \tag{15}$$

$$e(x, y) = \begin{cases} \sum_{d=1}^{\infty} p_d(\tau(x) - \tau(y)) e(y, z_1)^d e(y, z_2)^d + 1 - g_{\tau(x)-\tau(y)} & y \notin V_L(L) \\ 1 - g_{\tau(x)-\tau(y)} & y \in V_L(L) \end{cases} \tag{16}$$

$e(u) = e(u, w)$ where $w$ is the other child of $u$ if it were not lost. Equations from 11 to 16 are derived in [2].

### 2 Species Network Operators

Because we fix the topology of the species network, the operators in this group only modify the speciation and reticulation times in the species network. Also, to improve the computational performance of the MAP search, we calculate and utilize the Hastings ratios of operators in this group. Here we describe the species network operators.

**Scale-Time.** In this operator, we scale the speciation times $\tau$ of all internal vertices of the network $\mathbb{S}$ by a scale factor $r$ so that $\tau' = r\tau$. $r$ is drawn from Uniform$(f, \frac{1}{f})$ where $f$ is a small value close to 1. Moving between $(\tau, r)$ and $(\tau', r')$ means that $r' = \frac{1}{r}$, so the Hastings ratio is:

$$\frac{q(r')}{q(r)}\left|\frac{\partial(\tau', r')}{\partial(\tau, r)}\right| = \frac{1}{\frac{1}{f} - f} / \frac{1}{\frac{1}{f} - f}\left|\begin{matrix} \partial\tau'/\partial\tau & \partial\tau'/\partial r \\ \partial(1/r)/\partial\tau & \partial(1/r)/\partial r \end{matrix}\right| \tag{17}$$

$$= \left|\begin{matrix} r\mathbf{I} & \tau \\ 0 & r^{-2} \end{matrix}\right| = r^{|IV(S)|-2} \tag{18}$$

Due to the temporal constraint on coalescent times, which must be older than speciation times and younger than the duplication time of the corresponding locus, proposed species networks may be incompatible with the gene trees and be rejected on that basis.

**Change-Time.** This operator changes the time of only one randomly selected vertex, unlike **Scale-Time** which scales all vertices. So, effectively, $|IV(S)| = 1$, and the Hastings ratio is $r^{-1}$ where $r$ is the scale factor as in **Scale-Time**. As with **Scale-Time**, species networks proposed by this operator may be rejected due to temporal constraints.

**Scale-Time-All**. In this operator, the times $\tau$ of all internal vertices of $\mathbb{S}$, $\mathbb{L}$ and $\mathbb{G}$ are scaled by a factor $r$ and modified into $\tau' = r\tau$. $r$ is drawn from Uniform$(f, \frac{1}{f})$ where $f$ is a small value close to 1. Moving between $(\tau, r)$ and $(\tau', r')$ requires that $r' = \frac{1}{r}$, so the Hastings ratio is:

$$\frac{q(r')}{q(r)}\left|\frac{\partial(\tau', r')}{\partial(\tau, r)}\right| = \frac{1}{\frac{1}{f} - f} / \frac{1}{\frac{1}{f} - f}\left|\begin{matrix} \partial\tau'/\partial\tau & \partial\tau'/\partial r \\ \partial(1/r)/\partial\tau & \partial(1/r)/\partial r \end{matrix}\right| = \left|\begin{matrix} r\mathbf{I} & \tau \\ 0 & r^{-2} \end{matrix}\right| \tag{19}$$

$$= r^{|IV(S)|+|V_D(L)|+|IV(G)|-2} \tag{20}$$

**Change-Time-All.** A species network internal vertex $x$ with height $\tau(x)$ is selected at random. A random value $r$ is drawn from Uniform$(f, \frac{1}{f})$ where $f$ is a small value close to 1. $\tau$ is modified to a new value $\tau(x)' = r\tau(x)$. At the same time, the heights of associated locus network and gene tree vertices will also be modified. Assume that $x$ has parent(s) $y_1$ (and possibly $y_2$) and child (children) $z_1$ (and possibly $z_2$). Specifically, for $y_1$, assume that there is a set $D_1$ of duplication vertices reconciled to species network branch $(x, y_1)$ and a set $C_1$ of gene tree vertices reconciled to locus network branches with both vertices reconciled to either $x$ or $y_1$ or $(x, y_1)$. If there exists $y_2$, $D_2$ and $C_2$ are defined for $y_2$ similarly, if there doesn't exist $y_2$, $D_2 = \emptyset$ and $C_2 = \emptyset$.

Let $h_1 = \tau(y_1) - \tau(x)$ and $h_1' = \tau(y_1) - \tau(x)'$. The heights of $D_1$ and $C_1$ will be scaled. For vertex $u \in D_1 \cup C_1$, the new height will be: $\tau'(u) = \tau(y_1) - \frac{h_1'}{h_1}(\tau(y_1) - \tau(u))$. If $y_2$ is present, the same is done for $y_2$. Specifically, for $z_1$, assume that there are set $D_3$ of duplication vertices reconciled to species network branch $(x, z_1)$ and set $C_3$ of gene tree vertices reconciled to locus network branches with both vertices reconciled to either $x$ or $z_1$ or $(x, z_1)$. If there exists $z_2$, $D_4$ and $C_4$ are defined for $z_2$ similarly, if there doesn't exist $z_2$, $D_4 = \emptyset$ and $C_4 = \emptyset$.
Let $h_3 = \tau(x) - \tau(z_1)$ and $h_3' = \tau'(x) - \tau(z_1)$. The heights of $D_3$ and $C_3$ will be scaled. Assume for vertex $u \in D_3 \cup C_3$, the new height will be: $\tau'(u) = \tau(z_1) + \frac{h_3'}{h_3}(\tau(u) - \tau(z_1))$ If $z_2$ is present, the same is done for $u \in D_4 \cup C_4$. So the Hastings ratio is :

$$\frac{q(r')}{q(r)} \left| \frac{\partial(\tau', r')}{\partial(\tau, r)} \right| = \frac{1}{\frac{1}{f} - f} / \frac{1}{\frac{1}{f} - f} \left| \begin{matrix} \partial\tau'/\partial\tau & \partial\tau'/\partial r \\ \partial(1/r)/\partial\tau & \partial(1/r)/\partial r \end{matrix} \right| \tag{21}$$

$$= r^{-1} (\frac{h_1'}{h_1})^{(|D_1|+|C_1|)} (\frac{h_2'}{h_2})^{(|D_2|+|C_2|)} (\frac{h_3'}{h_3})^{(|D_3|+|C_3|)} (\frac{h_4'}{h_4})^{(|D_4|+|C_4|)} \tag{22}$$

### 3 Operators on the Locus Networks

We describe operators that change a locus network and also the gene tree.

**Shuffle-Reconciliation**. Because we model the reconciliation between the gene tree and locus network by reconciling coalescent events to the locus network branches, but integrate out the possible pathways for the edges leading to each coalescent event, Figure 1A and Figure 1B are two valid reconciliations for the same locus network and gene tree that must be sampled. In this operator, we shuffle the reconciliation by randomly choosing from one of the possible reconciliations.

**Move-Up-Down-Duplications**. In this operator, we define two complementary moves. One shifts the duplication event down to the child species, called **Dup-Move-Down** while the other shifts the duplications in the child species up into the mother species, denoted as **Dup-Move-Up**. With this operator, we are able to change the location of duplications (Figure 2). If the state is in Figure 2.a, i.e. there is only one duplication event $V_1$ mapped the the species network branch $(F, A)$ and $V_1$ has two children $A_1$ and $A_2$, where the corresponding gene tree lineages $a_1$ and $a_2$ does not coalesce in branch $(F_1, V_1)$, **Dup-Move-Up** can be applied to modify the state to Figure 2.b. We can move the duplication events up and map them to $(pa(F), F)$ to form $V_2$. Assume $l_1 = \tau(w_1) - \tau(F)$, a random number $r \sim U(0, 1)$ is generated, the height of the newly created $V_2$ is $\tau(V_2) = rl_1 + \tau(F)$. The gene tree is unchanged and $V_1$ will be deleted. If the locus network is in Figure 2.b, **Dup-Move-Down** can be used to restore to Figure 2.a. Assume the bound for adding the duplication is $l_2 = \tau(F) - \tau(w_3)$ where $w_3$ is the highest lower bound. In this case, $w_3$ can be $a_1$ or $a_2$, a random number $r \sim U(0, 1)$ is generated, the height of the newly created $V_1$ is $\tau(V_1) = rl_2 + \tau(w_3)$.

**Add-Remove-Single-Locus**. If a duplication vertex has one child, we call it a single locus ($pa(A_2)$ in Figure 3 (right)), it can be trivially removed or added back. The operator includes two moves **Remove-Single-Locus** and **Add-Single-Locus**. **Add-Single-Locus** is used to add single locus onto the locus network and the single locus can be deleted by **Remove-Single-Locus**.

**Add-Remove-Reti-Branch**. A branch $(u, v)$ is called a reticulate branch if $v$ is a reticulation vertex. We can add or remove reticulate branches with **Add-Reti-Branch** and **Remove-Reti-Branch** respectively (Figure 4). This is done because we assume that lineages can come only from one population leading to the reticulation vertex where the corresponding locus in the population of another parent is lost.

**Relocate-Duplications**. We can change a duplication time, constrained by upper and lower bounds. This operation will not cause the duplication vertex to be mapped to another species network branch. We denote the current height of the duplication as $\tau$. For a duplication vertex $u$, the upper bound $\tau_u$ is the minimum height among the speciation, duplication, or coalescent events above $u$. The lower bound $\tau_l$ is the maximum height among the speciation, duplication, or coalescent events below $u$ . We sample $\tau' \sim \text{Uniform}(\tau_l, \tau_u)$.

**Locus-Net-SPR**. This operator will cut and graft a locus to another position. For example, in Figure 5.a, the locus $B_2$ is the child of $V_1$, the coalescence event of $B_2$ is at $w_1$. $W_2$ is one potential location (among others) for relocating the locus. We can then cut off $B_2$ and reattach it with a new branch length to somewhere in the tree randomly (in this case $V_2$, see the path from $V_2$ to $B_2$). Because we allow single locus vertices on the locus network, our algorithm may sometimes add single locus vertices which results in $V_3$. The associated gene subtree defined as the child of the coalescent event $w_1$ above $B_2$ is simultaneously grafted to a compatible gene tree branch, creating a new coalescent event $w_2$, and reconciled to the locus tree branch $(pa(V_2), V_2)$.

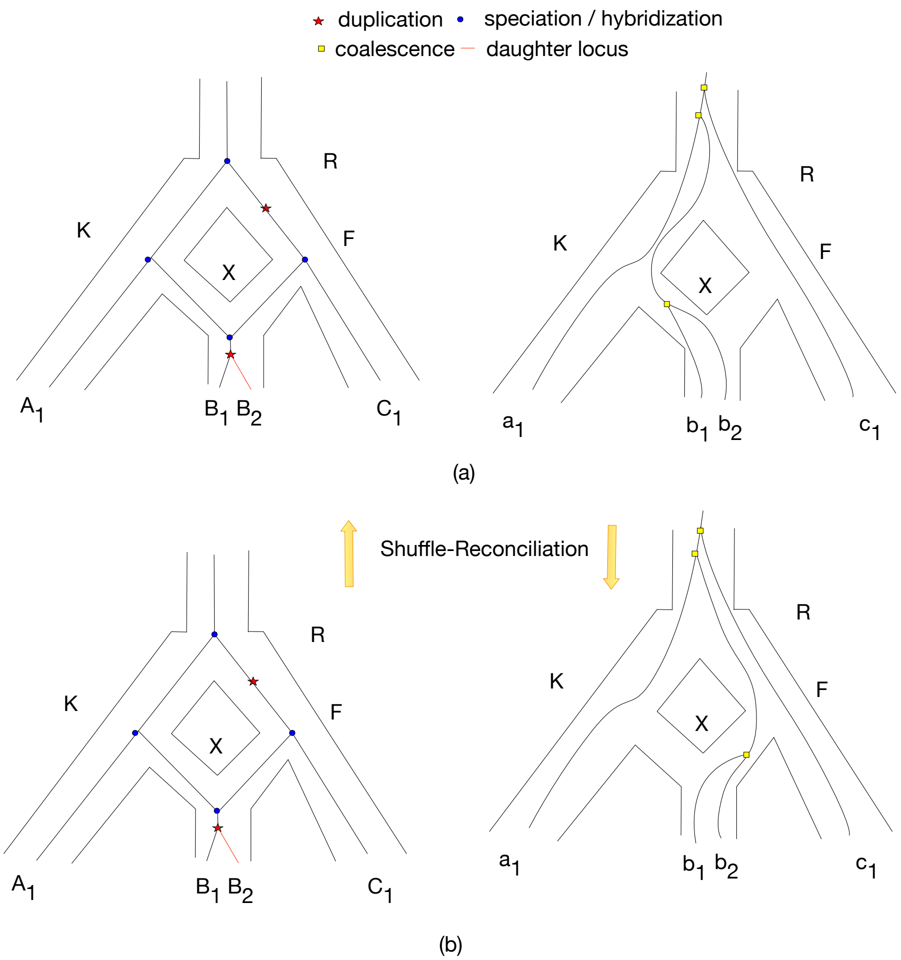


**Fig. 1.** The Shuffle-Reconciliation Operator. (a) and (b) are two valid reconciliations of the same locus network and gene tree. A shuffle operator can be applied to select randomly from one of them.

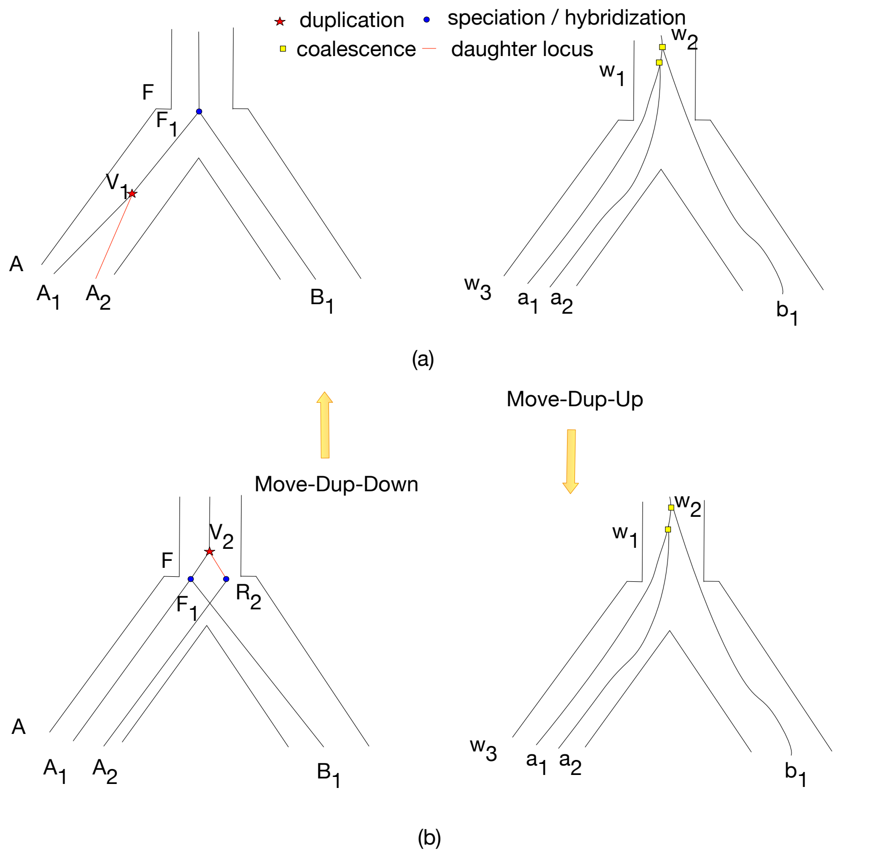


**Fig. 2.** Complementary moves made by the **Move-Up-Down-Duplications** operator. The duplication is on the $(F, A)$ giving 2 loci for species A in (a). It can be moved up to $(pa(F), F)$ to form (b). Also, if the state is in (b), the duplication can be moved down to be on $(F, A)$.

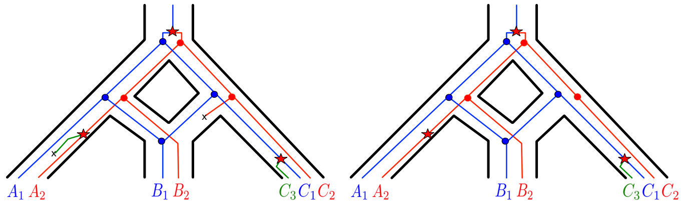


**Fig. 3.** A gene duplication and loss scenario inside of a species network on three species $A$, $B$, and $C$. (Left) The complete locus network embedded in the species network, produced by a birth-death process, and containing all duplication and loss events. (Right) Lineages in the locus network with no sampled loci due to loss events are pruned from the locus network, resulting in the observed locus network. Extinct lineages are deleted. Duplication, loss, and speciation/hybridization events are represented by $\star$, $\times$, and $\bullet$, respectively. New lineages arising from duplication are colored red and green.

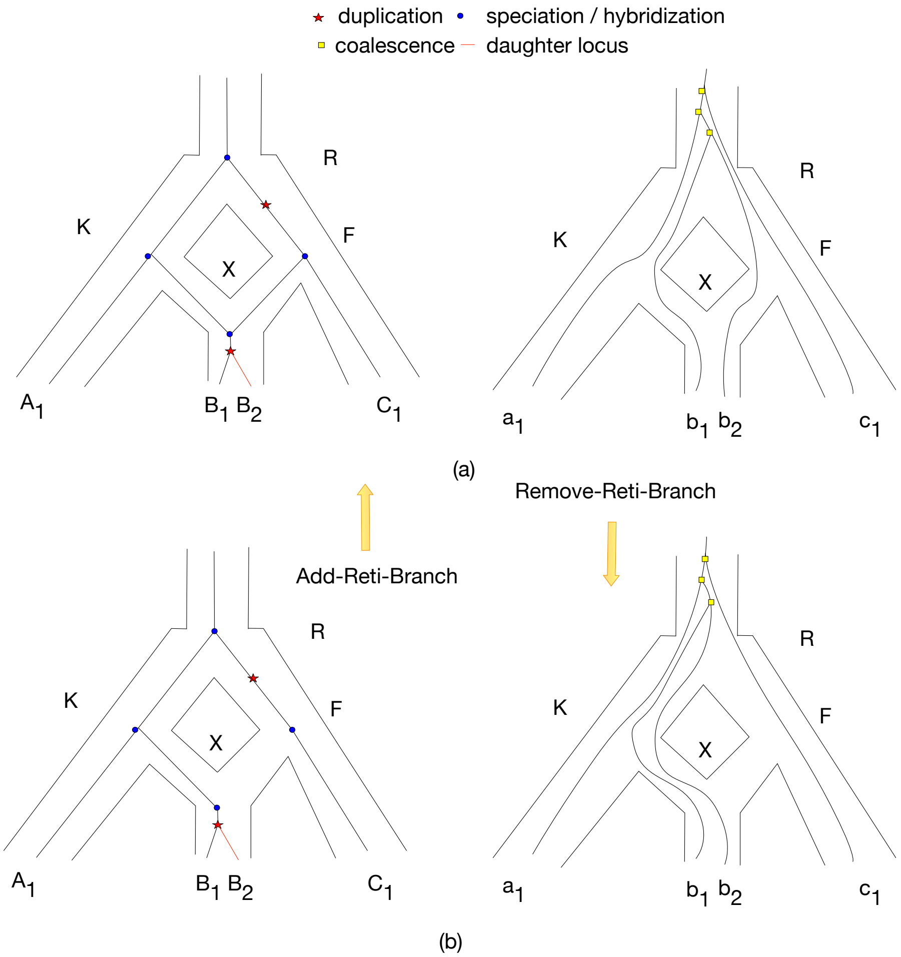


**Fig. 4.** The Add-Remove-Reti-Branch operator. This operator consists of two sub operators The reticulation branch embedded in (F, X) can be removed to generate (b) by operator **Remove-Reti-Branch** and restored to (a) by **Add-Reti-Branch**. **Remove-Reti-Branch** can only be done if doing so will not cause failure of reconciling gene tree in to the locus network.

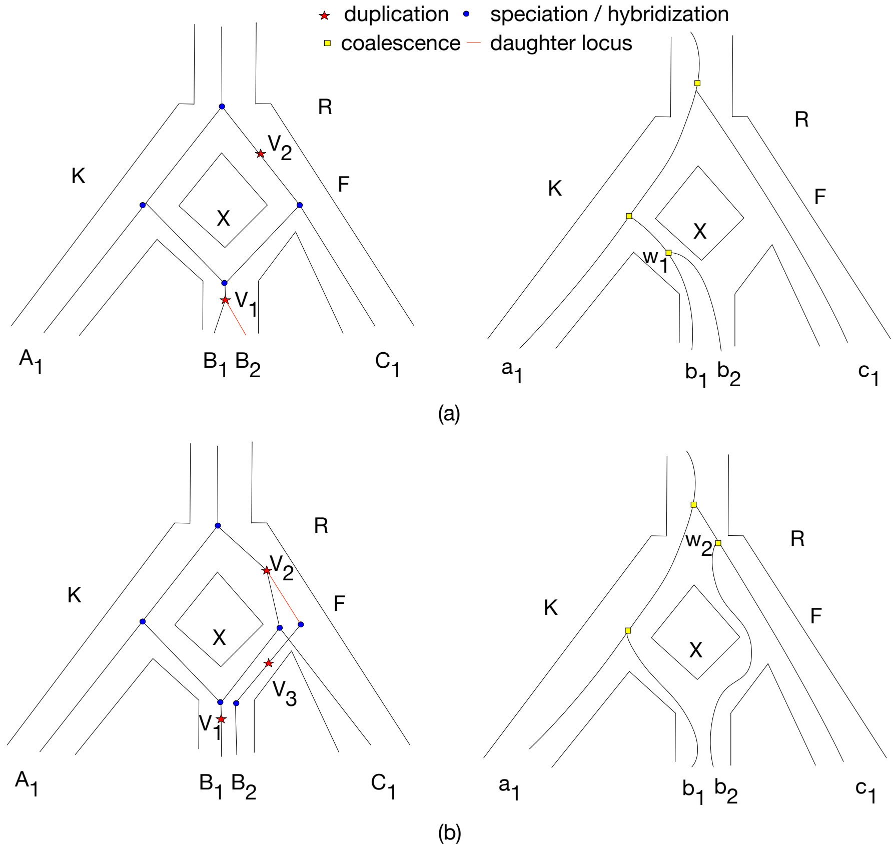


**Fig. 5.** The Locus-Net-SPR operator. Each leaf locus is attached a certain vertex in the locus network, we randomly picked one and then reattach it to another vertex. In this case, leaf locus $B_2$ is pruned off from $V_1$ and reattached to $V_2$ while the coalescence location is also changed.

### 4 Operators on the Gene Trees

**Scale-Time** In this operator, we scale the coalescent times $\tau$ of all internal vertices of the tree $\mathbb{G}$ by a scale factor $r$ and modify $\tau$ into $\tau' = r\tau$. $r$ is drawn from Uniform($f, \frac{1}{f}$) where $f$ is drawn uniformly from $(0, 1)$. As with the species network **Scale-Time** operator, species networks proposed by this operator may be rejected due to temporal constraints.

**Change-Time.** In this operation, the speciation times $\tau$ of one randomly selected vertex $u$ of the tree $\mathbb{G}$ are scaled by a scale factor $u$ and modified into $\tau' = r\tau$. $r$ is drawn from Uniform($f, \frac{1}{f}$) where $f$ is drawn uniformly from $(0, 1)$. As with the species network **Change-Time** operator, species networks proposed by this operator may be rejected due to temporal constraints.

**Gene-Tree-SPR**. In this operator, we will cut a subtree in the gene tree from its parent and reattach to a new node of the same height as the original parent on another branch. Subtrees will only be reattached to gene tree branches which are unambiguously mapped to the same locus network branch as the original subtree parent.

**Shuffle-Labeling**. In this operator, for a randomly selected gene family, we generate a new permutation of locus assignments at random for the gene copies that have been sampled within each species.

### 5 Operators on the Non-tree parameters

**Change-Dup-Rate.** The duplication rate $\mu$ is modified into $\mu'$ with the proposal:

$$\begin{cases} \mu' = \mu + u \text{ if } \mu + u \geq 0 \\ \mu' = -(\mu + u) \text{ if } \mu + u < 0 \end{cases} \tag{23}$$

where $u \sim$ Uniform$(-c, c)$ where c is a user defined small number.

**Change-Loss-Rate.** The loss rate $\lambda$ is modified into $\lambda'$ with the proposal:

$$\begin{cases} \lambda' = \lambda + u \text{ if } \lambda + u \geq 0 \\ \lambda' = -(\lambda + u) \text{ if } \lambda + u < 0 \end{cases} \tag{24}$$

where $u \sim$ Uniform$(-c, c)$ where c is a user defined small number.

### 6 Search Heuristic

We use a stochastic hill-climbing (SHC) algorithm for searching for the optimal point as given by

$$(\mathbb{S}^*, \mathbb{GF}^*, \theta^*) = \text{argmax}_{(\mathbb{S}, \mathbb{GF}, \theta)} p(\mathbb{S}, \mathbb{GF}, \theta | D). \tag{25}$$

and accept each proposed state according its unnormalized posterior probability $np$ relative to that of the current state $op$, similar to the Metropolis-Hastings algorithm [4]:

$$ac = min\left(1, w(\frac{np}{op})^{1/T}\right) \tag{26}$$

where $ac$ is the probability of accepting the proposed state and $w$ is a weight. Our optimization has two notable heuristics. First, we use a simulated healing algorithm [1] to flatten the posterior space, raising the acceptance probability and helping jump out of local optima for a prespecified fraction of iterations at the beginning of the chain, also we set the temperature $T$ hotter for locus network operators to further assist efficiency of those operators. Second, we use Hastings ratios when making changes on species network parameters, i.e. $w$ is set to the proposal ratio $P(s|s') \div P(s'|s)$ where $s$ and $s'$ are the current and proposed states.
